# rstoolbox: management and analysis of computationally designed structural ensembles

**DOI:** 10.1101/428045

**Authors:** Jaume Bonet, Zander Harteveld, Fabian Sesterhenn, Andreas Scheck, Bruno E. Correia

## Abstract

**Motivation:** Computational protein design (CPD) calculations rely on the generation of large amounts of data on the search for the best sequences. As such, CPD workflows generally include the batch generation of designed decoys (sampling) followed by ranking and filtering stages to select those with optimal metrics (scoring). Due to these factors, the proper analysis of the decoy population is a key element for the effective selection of designs for experimental validation.

**Results:** Here, we present a set of tools for the analysis of protein design ensembles. The tool is oriented towards protein designers with basic coding training aiming to process efficiently their decoy sets as well as for protocol developers interested in benchmarking their new approaches. Although initially devised to process Rosetta design outputs, the library is extendable to other design tools.

**Availability and Implementation:** rstoolbox is implemented for python2.7 and 3.5+. Code is freely available at https://github.com/lpdi-epfl/rstoolbox under the MIT license. Full documentation and examples can be found at https://lpdi-epfl.github.io/rstoolbox.

## 1 Introduction

Computational protein design (CPD) has become a popular tool for the creation and optimization of new proteins and functionalities (Gainza-Cirauqui and Correia 2018). Due to the abysmal size of the sequence-structure space (Taylor, Chelliah et al. 2009), CPD heuristic approaches became necessary. These approaches, such as Monte Carlo (MC) (Li and Scheraga 1987), use stochastic sampling methods and a scoring function to guide the structure and sequence exploration towards an optimal score. This allows them to explore a wider area of the sequence-structure space in a feasible time span. However, these approaches cannot guarantee that the solutions reached the global minima (Gainza, Nisonoff et al. 2016). CPD workflows relying in heuristic approaches compensate for this shortcoming in two way: I) extensive sampling yielding a large decoy population; II) sophisticated ranking and filtering schemes to aid the recognition of the best solutions. This general approach is used by the Rosetta suite (Alford, Leaver-Fay et al. 2017), one of the most widespread CPD tool.

For Rosetta, as with other similar approaches, the amount of sampling required scales with the degrees of freedom (conformational and sequence) that a particular CPD task demands. While deterministic routines such as scoring only require a single output, full structure prediction such as *ab initio* or docking may require to generate up to 10^6^ decoys to actually find acceptable solutions amongst them (Simons, Bonneau et al. 1999, Kim, Blum et al. 2009). Between these two extremes, a variety of sampling ranges have been estimated for other design problems. Thus, fixed backbone design simulations (Kuhlman and Baker 2000) may reach sufficient sampling within hundreds of decoys while protocols that allow backbone flexibility such as loop modeling (Stein and Kortemme 2013) or flexible backbone (Kuhlman, Dantas et al. 2003) design might go up to 10^4^ and 10^5^ decoys respectively. Due to the high number of decoys generated in the search of the best design solution, tools to facilitate the management and analysis of large decoy ensembles are essential to CPD pipelines.

Here, we present rstoolbox, a python library aimed to the management and analysis of design populations. The library includes scripts for quick data overview and presents a full set of functions to produce multi-parameter scoring schemes and compare design ensembles generated from different protocols. The full bundle of functionalities are useful and accessible to designers with limited coding experience, through visual interfaces such as Ipython (Pérez and Granger 2007), and also to developers interested in benchmarking and optimizing new CPD protocols.

## 2 Package Overview

The rstoolbox library has been written over pandas (McKinney 2010). Its main component is the DesignFrame, a table-like structure in which rows represent designed decoys and columns particular properties of the decoy (including score terms, sequence, secondary structure, residues of interest and others). This structure directly allows for key functionalities such as sequence variant identification, sequence similarity analysis or decoy selection and access to the specific plotting functions included in the library.

DesignFrames are easily filled by reading Rosetta silent files, a compressed format that holds both structure coordinates and statistics of Rosetta outputs. Additionally, any tabulated or table-like data file can be casted into a DesignFrame. This makes the library effortlessly adaptable to work with other design tools.

The components of the library can directly interact with most of the commonly used python plotting libraries. In addition, extra plotting functions are also present to facilitate specific analysis of CPD data. Although the library can access Rosetta functions to perform extra functionalities such as automatic backbone angle determination and Ramachandran analysis, most of its functionalities are independent of a local installation of Rosetta. This dramatically reduces the requirements and complexity for non-regular programmers to setup and exploit the library.

**Fig. 1.**
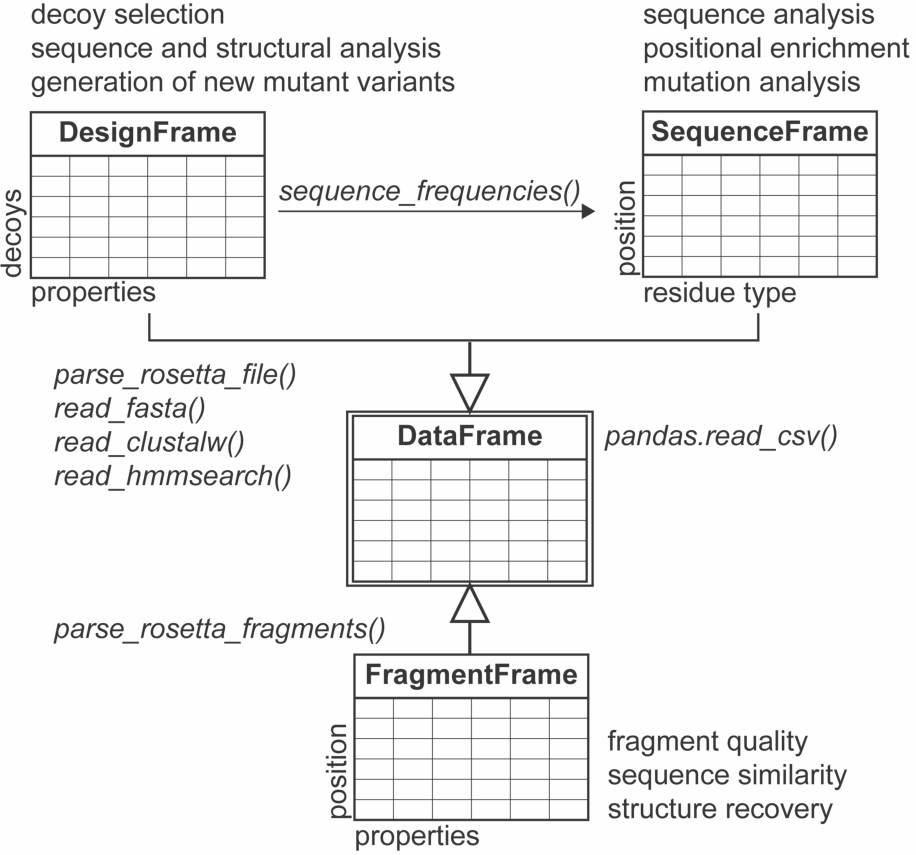
rstoolbox structure. The library relies on multiple table-like objects derived from DataFrames. Each object is created through different reading functions for different input types and provide access to a variety of analysis. Due to their table-like nature, tabulated formats are widely compatible, enabling the easy input and process new data formats.

## 3 Results

### 3.1 Direct Executables

The rstoolbox provides a set of direct executables described in the library documentation and exemplified in the Supplementary Materials. Of special note is minisilent.py, which removes all structural data from silent files reducing the file size dramatically. This becomes an extremely useful solution to include CPD metadata into public repositories, facilitating the management of published datasets and, thus, increasing reproducibility.

### 3.2 Library Access

The library exposes a set of functions (each of them explained in detail in the documentation API) that allows both for the rich analysis of a given dataset as well as the setup of new design simulations. On one hand, it provides visual tools (see Supplementary Material) and multi-score filtering as means of analysis. On the other hand, it can generate PSSM matrices and promote new mutants from selected decoys, producing the necessary files to explore those new mutants in Rosetta. Through this, it becomes the ideal solution for the human-guided steps between CPD cycles. Covering the final steps of the experimental testing of designs generated with CPD, the library can manage experimental data obtained properly attach it to the individual decoy. rstoolbox allows for computational and experimental data to be integrated, aiding the iterative process of experimental assessment and optimization coupled to CPD to generate improved designs.

## Availability

The rstoolbox source code, a complete API and documentation, examples, installation instructions, issue tracking and continuous integration are available via the GitHub repository. For users-only, the package is available at PyPI with the call ‘pip install rstoolbox’.

## References

Alford, R. F., et al. (2017). “The Rosetta All-Atom Energy Function for Macromolecular Modeling and Design.” J Chem Theory Comput 13(6): 3031–3048.

Gainza, P., et al. (2016). “Algorithms for protein design.” Curr Opin Struct Biol 39: 16–26.

Gainza-Cirauqui, P. and B. E. Correia (2018). “Computational protein design-the next generation tool to expand synthetic biology applications.” Curr Opin Biotechnol 52: 145–152.

Kim, D. E., et al. (2009). “Sampling bottlenecks in de novo protein structure prediction.” J Mol Biol 393(1): 249–260.

Kuhlman, B. and D. Baker (2000). “Native protein sequences are close to optimal for their structures.” Proc Natl Acad Sci U S A 97(19): 10383–10388.

Kuhlman, B., et al. (2003). “Design of a novel globular protein fold with atomic-level accuracy.” Science 302(5649): 1364–1368.

Li, Z. and H. A. Scheraga (1987). “Monte Carlo-minimization approach to the multiple-minima problem in protein folding.” Proc Natl Acad Sci U S A 84(19): 6611–6615.

McKinney, W. (2010). “Data Structures for Statistical Computing in Python.” Proceedings of the 9th Python in Science Conference: 51–56

Pérez, F. and E. B. Granger (2007). “IPython: A System for Interactive Scientific Computing.” Computing in Science and Engineering 9(3): 21–29.

Simons, K. T., et al. (1999). “Ab initio protein structure prediction of CASP III targets using ROSETTA.” Proteins **Suppl** 3: 171–176.

Stein, A. and T. Kortemme (2013). “Improvements to robotics-inspired conformational sampling in rosetta.” PLoS One 8(5): e63090.

Taylor, W. R., et al. (2009). “Probing the “dark matter” of protein fold space.” Structure 17(9): 1244–1252.

